# A temperature-sensitive *Mycobacterium smegmatis glgE* mutation leads to a loss of GlgE enzyme activity and thermostability and the accumulation of α-maltose-1-phosphate

**DOI:** 10.1101/2020.04.23.057067

**Authors:** Karl Syson, Sibyl F. D. Batey, Steffen Schindler, Rainer Kalscheuer, Stephen Bornemann

## Abstract

**Background:** The bacterial GlgE pathway is the third known route to glycogen and is the only one present in mycobacteria. It contributes to the virulence of *Mycobacterium tuberculosis*. The involvement of GlgE in glycogen biosynthesis was discovered twenty years ago when the phenotype of a temperature-sensitive *Mycobacterium smegmatis* mutation was rescued by the *glgE* gene. The evidence at the time suggested *glgE* coded for a glucanase responsible for the hydrolysis of glycogen, in stark contrast with recent evidence showing GlgE to be a polymerase responsible for its biosynthesis.

**Methods:** We reconstructed and examined the temperature-sensitive mutant and characterised the mutated GlgE enzyme.

**Results:** The mutant strain accumulated the substrate for GlgE, α-maltose-1-phosphate, at the non-permissive temperature. The glycogen assay used in the original study was shown to give a false positive result with α-maltose-1-phosphate. The accumulation of α-maltose-1-phosphate was due to the lowering of the *k*_cat_ of GlgE as well as a loss of stability 42 ºC. The reported rescue of the phenotype by GarA could potentially involve an interaction with GlgE, but none was detected.

**Conclusions:** We have been able to reconcile apparently contradictory observations and shed light on the basis for the phenotype of the temperature-sensitive mutation.

**General Significance:** This study highlights how the lowering of flux through the GlgE pathway can slow the growth mycobacteria.

## 1. Introduction

A third biosynthetic pathway to produce glycogen has been elucidated relatively recently [1,2]. This involves the enzyme GlgE (Fig. 1) acting as the polymerase [1,3]. GlgE was first identified as an enzyme involved in glycogen metabolism in 1999 [4]. Intriguingly, it was then reported that a temperature-sensitive mutant appeared to accumulate glycogen, which is the opposite of what would be expected given our current understanding of the GlgE pathway. This study addresses this apparent discrepancy.

**Fig. 1.**
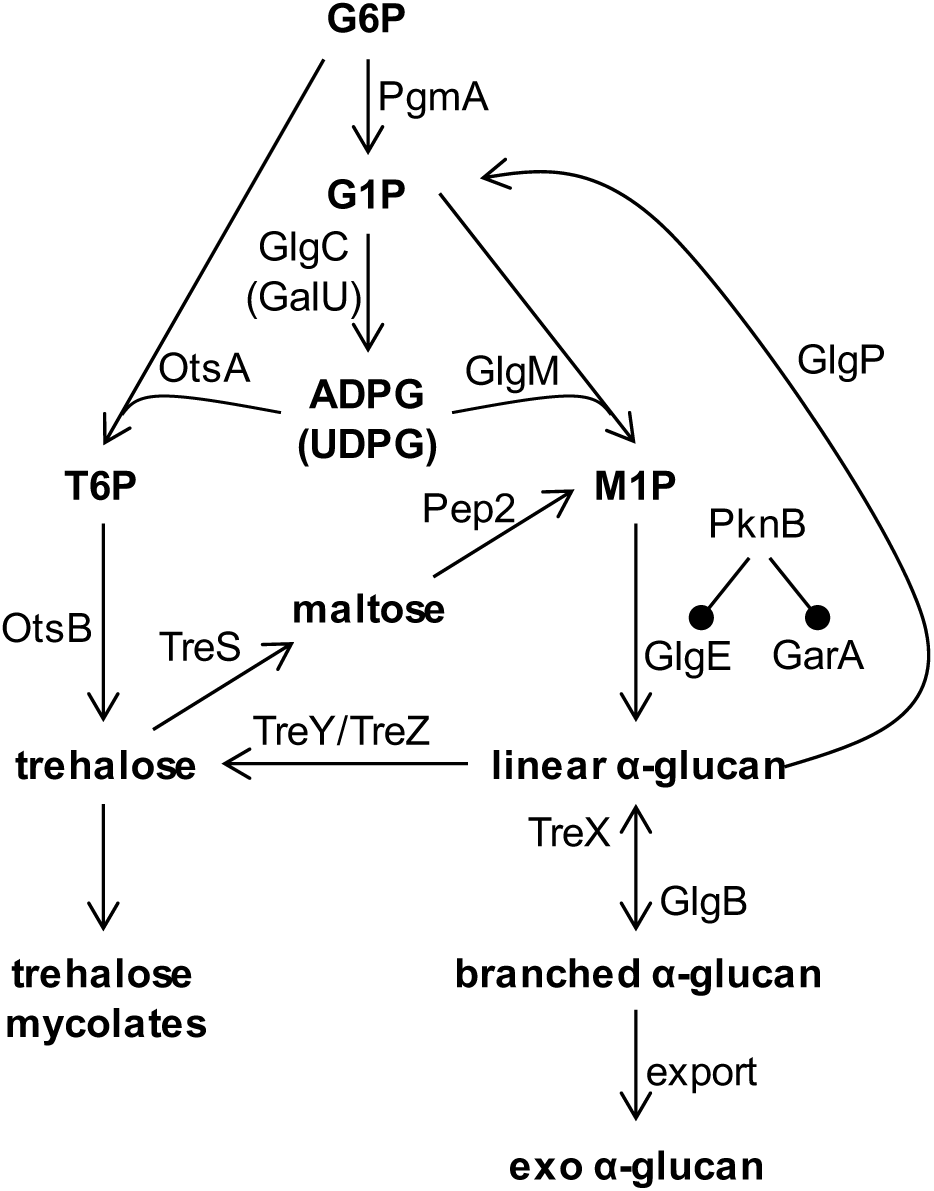
Glycogen (α-glucan) pathways in mycobacteria. The scheme shows how glucose-6-phosphate (G6P) can be converted to α-glucan *via* the following intermediates; glucose-1-phosphate (G1P), ADP-glucose (ADPG) or UDP-glucose (UDPG), trehalose-6-phosphate (T6P), and α-maltose-1-phosphate (M1P). The regulation of GlgE and GarA by PknB is indicated. The figure is adapted from [5].

Glycogen is well known as a carbon and energy storage molecule [6]. When a temperature-sensitive *Mycobacterium smegmatis* mutant that slowed growth and appeared to accumulate glycogen was isolated (SMEG53), the phenotype was complemented by the *glgE* gene [4]. The sequence of the genomic copy of the *glgE* gene revealed a His349Tyr amino acid substitution was present in the GlgE protein of the temperature-sensitive mutant. There appeared to be a correlation between the accumulation of glycogen with the cessation of exponential growth at the non-permissive temperature of 42 °C. Furthermore, there also appeared to be a correlation between the accumulation of glycogen and growth rate. Suppression of the temperature-sensitive phenotype was observed in certain growth conditions and by the over-expression of *garA*, which codes for a forkhead-associated domain protein consequently referred to as a glycogen accumulation regulator [7].

GlgE is a GH13_3 CAZy family member [8]. The GH13 family comprises proteins with several enzyme activities, most notably α-amylases [9]. On this basis, GlgE was initially predicted to be a hydrolase capable of degrading the α-1,4 linkages of glycogen (glucanase) [4]. Given the interpretation of the results of the above study and the bioinformatic prediction, it was concluded that GlgE is a glucanase and that it is involved in glycogen recycling that is essential for exponential growth.

However, it has subsequently and unequivocally been shown that GlgE possesses α-maltose-1-phosphate:(1→4)-α-D-glucan 4-α-D-maltosyltransferase polymerase-type activity (EC 2.4.99.16) and some disproportionation activity, but no detectable hydrolase activity [1,10]. The polymerases for each of the known glycogen biosynthetic pathways extend existing oligomers and polymers to generate linear α-1,4-linked chains [3,11]. In all three pathways, a branching enzyme GlgB then introduces α-1,6 branchpoints. The GlgE pathway uses α-maltose-1-phosphate as the building block for the biosynthesis of an α-glucan that is similar to classical glycogen except for its fine structure [3]. There are two routes to the formation of α-maltose-1-phosphate in mycobacteria (Fig. 1). One is via trehalose and maltose plus ATP, involving trehalose synthase and maltose kinase [1,2], while the other is via glucose-1-phosphate plus ADP-glucose involving α-maltose-1-phosphate synthase [5,12]. Importantly, the loss of the *glgE* gene in *M. smegmatis* is known to lead to the accumulation of α-maltose-1-phosphate and the slowing of bacterial growth [1,5]. The GlgE-derived glycogen is partially exported by *Mycobacterium tuberculosis* to form a loosely attached capsule that is associated with increased virulence [5,13]. Notably, the two classical glycogen biosynthetic pathways that use ADP-glucose or UDP-glucose [6] are now known to be absent in mycobacteria [5].

To address an apparent inconsistency between the initial study suggesting GlgE is a hydrolase and subsequent work showing that it is a polymerase, several hypotheses were tested. Since GlgE is a polymerase that uses α-maltose-1-phosphate, a variant of GlgE that is somehow compromised would be expected to accumulate this building block leading to a lower bacterial rate of growth, as has been observed in other studies [1]. It is therefore possible that the glycogen assay used in the 1999 study, based on an amyloglucosidase and the quantification of released glucose [4], was also capable of detecting α-maltose-1-phosphate. Interestingly, the GarA protein is known to regulate carbon and nitrogen metabolism through binding to other proteins [7]. Perhaps GarA binds to GlgE and rescues the temperature-sensitive version. However, it is also a substrate of the regulatory Ser/Thr protein kinase PknB, which in turn has also been shown to negatively regulate GlgE [14]. Perhaps GarA therefore rescues the GlgE mutation through interfering with the regulatory function of PknB. The present study aims to test these hypotheses and resolve the apparent inconsistencies. This was done by reconstructing and phenotyping the mutant strain together with testing the hydrolysis of α-maltose-1-phosphate by amyloglucosidase and characterising the H349Y variant of the GlgE enzyme.

## 2. Materials and methods

### 2.1. Recombinant M. smegmatis GlgE and GarA

The *M. smegmatis glgE* gene was synthesized with optimized codon usage for expression in *Escherichia coli* (Genscript Corporation, Piscataway, NJ), allowing the production of GlgE with a 6×His tag and a TEV cleavage site at its *N*-terminus. The gene coding for the H349Y GlgE variant was generated from the synthesized wild-type *glgE* gene using a QuikChange Lightening kit (Agilent). The constructs were ligated into pET21a expression vectors (Novagen, Darmstadt, Germany) using *Nde*I and *Bam*HI restriction sites. *E. coli* BL21 (DE3) Star (Novagen) bearing each plasmid were grown at 25 °C to an optical density of 0.6 at 600 nm in Lysogeny Broth (LB) and expression was induced with 0.5 mM isopropyl β-D-thiogalactopyranoside (IPTG). Bacteria were harvested and lyzed after a further 16 h incubation. The enzyme was purified using a 1 ml HisTrap FF column (GE Healthcare, Amersham, United Kingdom) with imidazole gradient elution and an S200 16/60 gel filtration column (Pharmacia Biotech, Amersham, United Kingdom) eluted with 20 mM Tris buffer, pH 8.5, containing 100 mM NaCl. GlgE-containing fractions were pooled and concentrated to ~1.5 mg/ml and aliquots were stored at −80 °C. An *M. smegmatis garA* construct was ligated into a pETPhos expression vector using *NdeI* and *BamHI* restriction sites [14]. *E. coli* BL21 (DE3) Star (Novagen) bearing this plasmid was grown at 30 °C to an optical density of 0.6 at 600 nm in LB and expression was induced with 0.2 mM IPTG. Bacteria were harvested and lyzed after a further 3 h incubation. The enzyme was purified using a 1 ml HisTrap FF column (GE Healthcare, Amersham, United Kingdom). The enzyme was dialysed into 20 mM Bis-Tris propane, pH 7.0, containing 150 mM NaCl and concentrated to 9 mg/ml and aliquots were stored at −80 °C.

### 2.2. Enzyme assays

Unless otherwise stated, all enzyme assays were carried out in 100 mM Bis-Tris propane, pH 7.0, containing 50 mM NaCl. GlgE activity was monitored at 21 °C using an end-point assay monitoring the quantitative release of inorganic phosphate with malachite green [15]. The concentration of free inorganic phosphate was estimated from a standard curve. α-Maltose-1-phosphate was synthesised as described earlier [10]. Reaction mixtures of 25 μl comprised 1 mM maltohexaose and 0.25 mM α-maltose-1-phosphate. Enzyme concentrations were such to allow reactions to progress linearly for 10 min with total donor consumption being < 40%. Reactions were quenched with 175 μl of malachite green and incubated for 20 min at 21 °C before the absorbance at 630 nm was measured on a SpectraMAX Plus microplate spectrophotometer using SOFTmax PRO 3.1.1 software. The effect of pH was determined using 100 mM MES (pH 6.0), Bis-Tris (pH 6.5), Bis-Tris propane (pH 7.0), HEPES (pH 7.5) and Tris (pH 8.0) buffers. Acceptor preference used maltose, maltotriose, maltotetroase, maltopentaose, maltohexaose and maltoheptaose at 1 mM each. The effect of salt was determined from 0 to 350 mM NaCl. Temperature de-activation of GlgE and GlgE H349Y was performed by pre-incubating protein at 21, 35, 40, 45 and 50 °C for 20 min and assaying at 21 °C. Temperature de-activation rescue experiments were performed as above, but in the presence of 50 µM *M. smegmatis* GarA. The effect of temperature on enzyme activity was determined by measuring initial rates (*v*0/[E]) over a temperature range of 25 to 55 °C in 5 °C increments, with reaction mixtures of 40 µl comprising 1 mM maltohexaose and 1 mM α-maltose-1-phosphate. Enzyme kinetics at permissive (30 °C) and non-permissive (42 °C) temperatures were performed with maltohexoase concentrations between 1 and 150 mM and 1 mM α-maltose-1-phosphate.

Initial rates were measured by quenching 6 μl reaction aliquots in 94 μl of 1 M HCl at time points from 1 to 8 min. The quenched reactions were incubated with 700 μl of malachite green assay solution for 20 min at 21 °C, and the absorbance at 630 nm was measured on a Perkin Elmer Lambda 18 or 25 spectrophotometer. Reaction rates were linear over at least the first 4 min. Enzyme kinetics for maltohexaose were fitted to a substrate inhibition model, as defined by equation, 1 using GraphPad Prism v8.

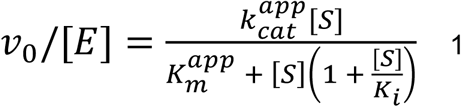

Equation 1 is a simplified form derived of equation 2 [16], where A would be α-maltose-1-phosphate and B the acceptor substrate in the case of GlgE [17].

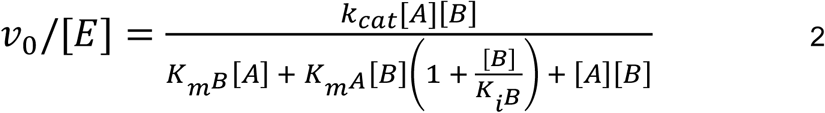

Equation 2 can rearranged to give equation 3.

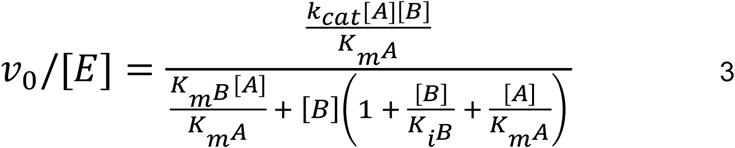

When the initial concentration of A (α-maltose-1-phosphate) is fixed and the initial concentration of B (acceptor) is varied, equation 3 appears in the form of equation 1 such that *k*_cat_^app^ and *K*_m_^app^ are defined by equations 4 and 5, respectively.

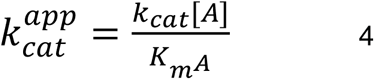

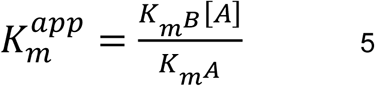

The hydrolysis of α-maltose-1-phosphate was tested using *Aspergillus niger* amyloglucosidase from Megazyme (high purity for use in Megazyme Total Starch method) in 100 mM sodium acetate buffer, pH 4.0, at 25 °C.

### 2.3. Thermofluor assay

Assay mixtures of 50 µl containing 0.6 mg/ml protein, 20 mM Bis-Tris propane, pH 7.0 and SYPRO Orange protein stain dye (Sigma). Melting curves were recorded with a temperature range of 20 to 95 °C, using a DNA Engine Opticon 3 real time PCR system (MJ Research) with Opticon Monitor 3.1 analysis software (BioRad).

### 2.4. Circular dichroism

Experiments were performed using a Chirascan-Plus CD spectrophotometer (Applied Photo-physics). Purified proteins were dialysed into 10 mM sodium phosphate, pH 7.0. CD measurements were carried out in a quartz glass cell with a 0.5 mm path length at a ~0.3 mg/ml protein concentration. To obtain overall CD spectra, wave-length scans between 180 and 260 nm were collected at 20, 30 and 42 °C using a 1.0 nm bandwidth, a 0.5 nm step size, and a time per point of 1 second. Triplicate spectra were averaged, followed by the subtraction of buffer only control spectra. Time course spectra were recorded when the sample reached 42 °C and again two hours later. The raw data in millidegree units were corrected for background and converted to mean residue molar ellipticity. Secondary structure assignments were made using the DichroWeb server [18], using the CDSSTR algorithm and reference set 7 [19]. Thermal melt curves were determined with wavelength scans between 195 and 260 nm using 1 nm bandwidth, 1.0 nm step size and time per point of 0.7 seconds. The temperature was increased from 20 to 90 °C at a rate of 1 °C per min and a complete scan was collected at 1 °C intervals. The raw data, in millidegree units, from 201 to 260 nm, at each temperature measured, were analyzed using the Global 3 software package (Applied Photophysics). The Global 3 package fits the full-spectrum data using the nonlinear regression method of Marquardt and Levenberg [20] to generate a global analysis of unfolding. This procedure determines fitted and optimized temperatures of transition (melting points: *T*_m_) and their associated Van’t Hoff enthalpies (Δ*H*).

### 2.5. Protein-protein interaction studies

Surface plasmon resonance experiments were carried out on a Biacore T200 system (GE Healthcare). Immobilisation of either GarA or GlgE was attempted using both amine coupling (CM5 chip) and nickel affinity (NTA chip) approaches as per the manufacturer’s instructions.

Fluorescence was recorded on a Perkin-Elmer LS55 fluorescence spectrometer connected to a PTP1 Peltier system. All assays were recorded using 100 mM Bis-Tris propane, pH 7.0, containing 50 mM NaCl. Anisotropy experiments used GlgE and GlgE variants labelled with dansyl chloride in 50 mM HEPES, pH 9.0, containing 50 mM NaCl over 45 mins at 21 °C [21], in a manner used by us in other systems [22]. Labelled protein was then dialysed into 20 mM Tris, pH 8.5. Excitation and emission wavelengths were 335 and 515 nm, respectively. Experiments were carried out by titrating GarA into 1 µM labelled protein, up to a concentration of 630 µM at 25 and 42 °C. Separately, tryptophan fluorescence was measured with 1 µM H349Y and a GarA concentration up to 100 µM.

### 2.6. Bacterial strains, growth curves and metabolite analyses

Two strains of *M. smegmatis* mc^2^155 were used. The parent strain was c-*glgE*-tet-off [5], which allowed the *glgE* gene to be silenced in the presence of anhydrotetracycline (ATc). A plasmid bearing the H349Y variant of the *glgE* gene, pMV361(Apra)::*glgE*-Ts, was introduced into this strain. Cells were grown aerobically at 30 or 42 °C in Middlebrook 7H9 medium supplemented with 0.5% v/v glycerol and 0.05% v/v tyloxapol and containing 10% v/v ADS enrichment (5% w/v bovine serum albumin fraction V (BSA), 2% w/v glucose, 0.85% w/v sodium chloride). Hygromycin (50 mg/l), kanamycin (20 mg/l) and apramycin (10 mg/l) were added for the selection of appropriate strains. Starter cultures of *M. smegmatis* strains with the appropriate antibiotic selection were grown at 30 °C in the presence or absence of ATc until they reached logarithmic phase. They were then diluted to an optical density of 0.3 at 600 nm and divided into two equal aliquots, one of which was grown at 30 °C and the other at 42 °C. When one of the cultures reached an optical density of ≥ 1.5 at 600 nm, they were each diluted to an OD of 1.5 (equating to 4.5 108 colony forming units per ml) and harvested by centrifugation, washed twice with phosphate buffered saline and resuspended in 2 ml of water. Soluble metabolites were released from cells by heating at 95 °C for 4 h, and cell debris was removed by centrifugation. The resulting supernatant was filtered, freeze-dried and re-suspended in 500 µl of D2O. ^1^H nuclear magnetic resonance spectra were recorded on a Bruker Avance III 400 MHz spectrometer at 22 °C and data were analysed using Topspin 3.5 software (Bruker Biospin Ltd). NMR signals were quantified by comparing integrals with sodium 3-(trimethylsilyl)propionate-2,2,3,3-d_4_ as an internal standard, which was also consistent with a standard curve generated with authentic trehalose.

## 3. Results

### 3.1. The temperature sensitive mutant accumulates α-maltose-1-phosphate at the non-permissive temperature

If the H349Y mutation in GlgE compromises enzyme activity, it would be expected that this would lead to the accumulation of α-maltose-1-phosphate [1]. To determine which small water-soluble metabolites accumulate in *M. smegmatis* as a result of the presence of the H349Y variant of GlgE at permissive and non-permissive temperatures, a gene silencing approach was used. The *glgE* gene has previously been brought under the control of a repressible promotor [5], allowing it to be robustly silenced in the presence of ATc. This was important because mutations in *glgE* are known to cause genetic instability over time because of toxicity associated with the accumulation of α-maltose-1-phosphate, particularly during growth in the presence of this metabolite’s precursor trehalose. To generate the temperature sensitive mutant, the H349Y variant of the *glgE* was introduced on a plasmid into the conditionally silenced strain. The addition of ATc to the growth medium would silence the wild-type copy making the H349Y variant the only GlgE present. Thus, it was possible to generate the temperature sensitive mutant synthetically, avoiding the possibility of additional mutations being introduced elsewhere in the genome during storage or manipulation. This would allow any phenotypes to be attributable to the single amino acid substitution in GlgE with confidence.

The two bacterial strains were then grown in the presence or absence of ATc at either the permissive or non-permissive temperatures of 30 and 42 °C, respectively. Growth kinetics confirmed that the c-glgE-tet-off strain expressing the H349Y variant of the *glgE* gene in the presence of ATc (Fig. 2) faithfully reproduced the temperature-sensitive growth phenotype reported for the SMEG53 strain [4]. This shows that the H349Y mutation of the *glgE* gene is sufficient to result in temperature-sensitivity at the cellular level.

**Fig. 2.**
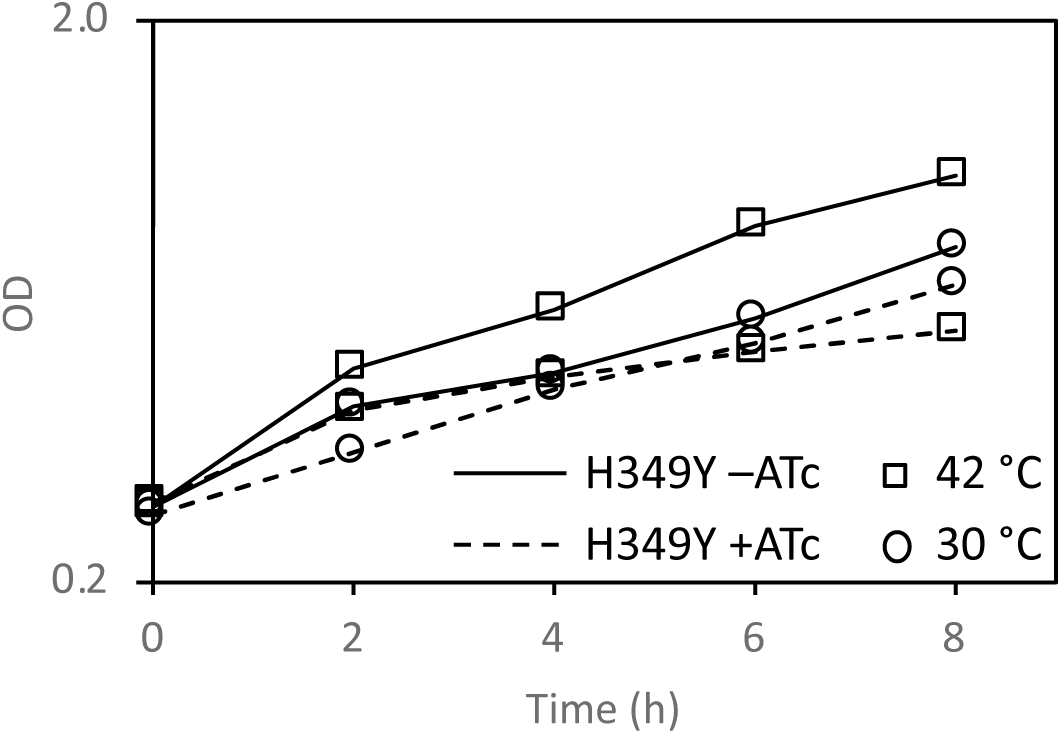
The growth of *M. smegmatis* is compromised by the H349Y mutation of GlgE, particularly at 42 °C. The optical density of cultures was monitored at 600 nm in two independent experiments and the mean values were plotted on a log scale. The c-*glgE*-tet-off pMV361(Apra)::*glgE-*Ts strain grew more rapidly at 42 than 30 °C in the absence of ATc (annotated as H349Y –ATC), as expected for a wild-type-like strain with a functional GlgE. This strain in the presence of ATc (annotated as H349Y +ATC) grew more slowly than the wild-type strain at each respective temperature. The temperature-sensitive strain initially grew more rapidly at 42 °C than at 30 °C, but growth was arrested after 2 hours of growth. These data with the reconstructed mutant faithfully recapitulate those reported for the isolated MSEG53 mutant [4].

Other phenotypes associated with SMEG53, such as altered colony morphology and partial rescue by osmolytes [4], were not observed in the reconstructed mutant strain in the presence of ATc. Since SMEG53 was isolated after treatment with the mutagen nitrosoguanidine and mutations in *glgE* are known to lead to genetic instability, it is likely that mutations in genes other than *glgE* might have contributed to the other phenotypes.

The soluble metabolites were then extracted from the strains grown under the different conditions and analysed using ^1^H-NMR spectroscopy. Whenever ATc was absent, the wild-type GlgE would be expected to be present. It would therefore be expected that no α-maltose-1-phosphate would accumulate [1] at any temperature, whether or not the H349Y variant of GlgE was present. This was indeed the case (Table 1). The only other small soluble carbohydrate metabolite that accumulated was trehalose, which was present in all strains including the wild-type, as reported previously. Interestingly, the level of trehalose increased at the higher temperature (e.g. from typically 29 to 46 µg/109 colony forming units). When ATc was added in the absence of the H349Y variant, no GlgE would be expected to be present. This would lead to the accumulation of α-maltose-1-phosphate (to non-lethal levels because no trehalose was added to the grown medium [1]). Accumulation indeed occurred, particularly at the higher temperature when more of the precursor, trehalose, was available. Finally, when ATc was added in the presence of the H349 variant, the mutated H349Y version of GlgE would be expected to be the only form present. When this strain was grown at 30 °C, no significant amount of α-maltose-1-phosphate was detected. However, a moderate amount of α-maltose-1-phosphate had clearly accumulated at 42 °C. One can therefore conclude that the temperature sensitivity of the H349Y GlgE strain led to the accumulation of α-maltose-1-phosphate implying GlgE was compromised by the mutation at the higher temperature, but not completely absent.

**Table 1.**
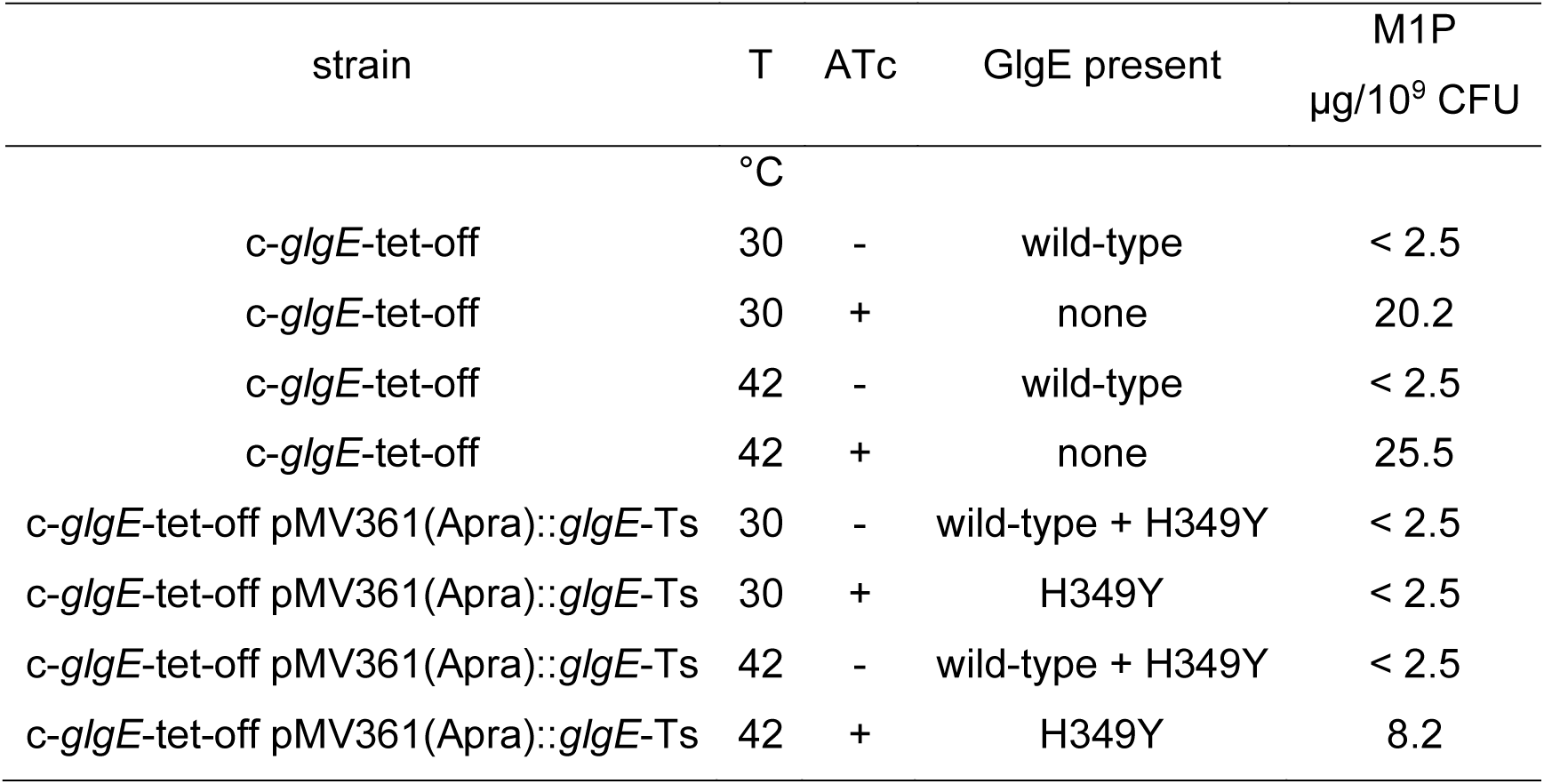
The temperature sensitive strain accumulates α-maltose-1-phosphate at the non-permissive temperature. The *glgE* gene in *M. smegmatis* was silenced with ATc using a tet-off system. The temperature sensitive H349Y variant of *glgE* was introduced into the parent strain on a plasmid. Each of the two strains were grown at either 30 or 42 °C in either the absence or presence of ATc. The expected presence of each variant of the GlgE enzyme is indicated in each case. The level of α-maltose-1-phosphate (M1P) was determined using ^1^H-NMR spectroscopy. The values represent means of two independent experiments (with a limit of detection of 2.5 µg/10^9^ colony forming units (CFU)). See also Fig. 3.

**Fig. 3.**
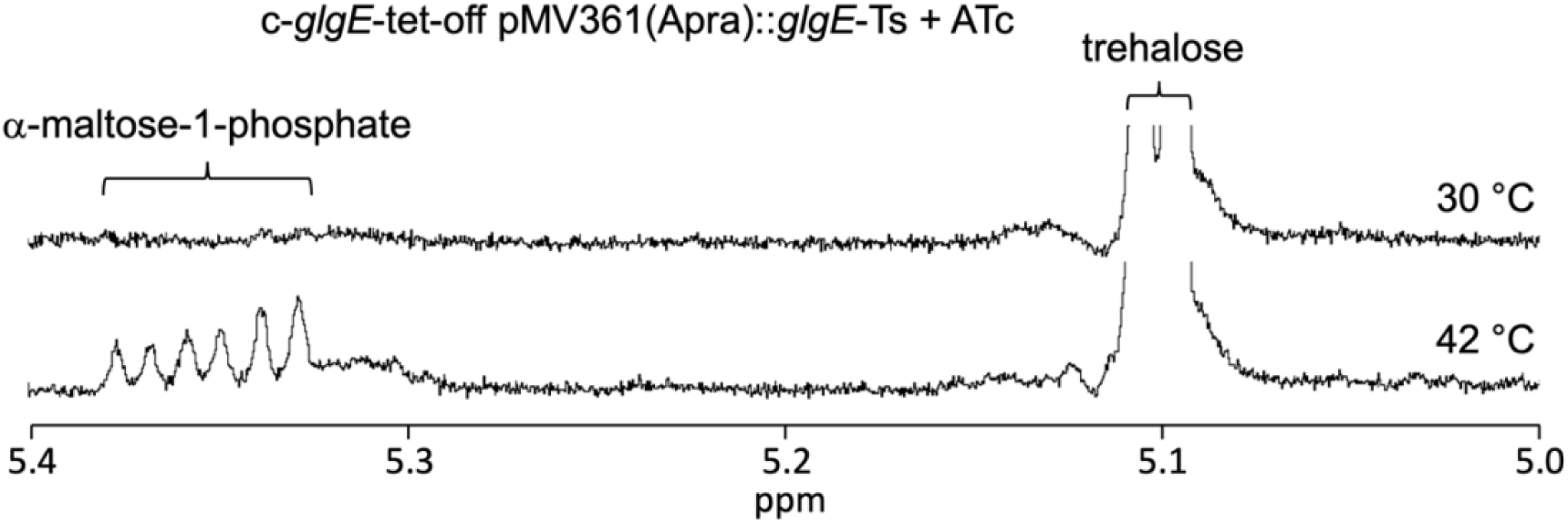
The H349Y mutation of GlgE leads to the accumulation of α-maltose-1-phosphate when *M. smegmatis* is grown at 42 °C. Representative ^1^H-NMR spectra of two biological repicates are shown of extracts of the c-*glgE*-tet-off pMV361(Apra)::*glgE-*Ts strain grown in the presence of ATc at 30 or 42 °C. See also Table 1.

### 3.2. α-Maltose-1-phosphate is hydrolysed by amyloglucosidase

An anomeric mixture of α*/*β-maltose-1-phosphate was incubated with *A. niger* amyloglucosidase. Both anomers were efficiently degraded by the enzyme to yield glucose-1-phosphate and glucose (Fig. 4). As expected for this type of enzyme (EC 3.2.1.3), the β anomer of glucose was liberated initially, which subsequently reaches an equilibrium with the α anomer within an hour or so. It is therefore clear that any α-maltose-1-phosphate that accumulates in *M. smegmatis* would be degraded by this enzyme to yield glucose.

**Fig. 4.**
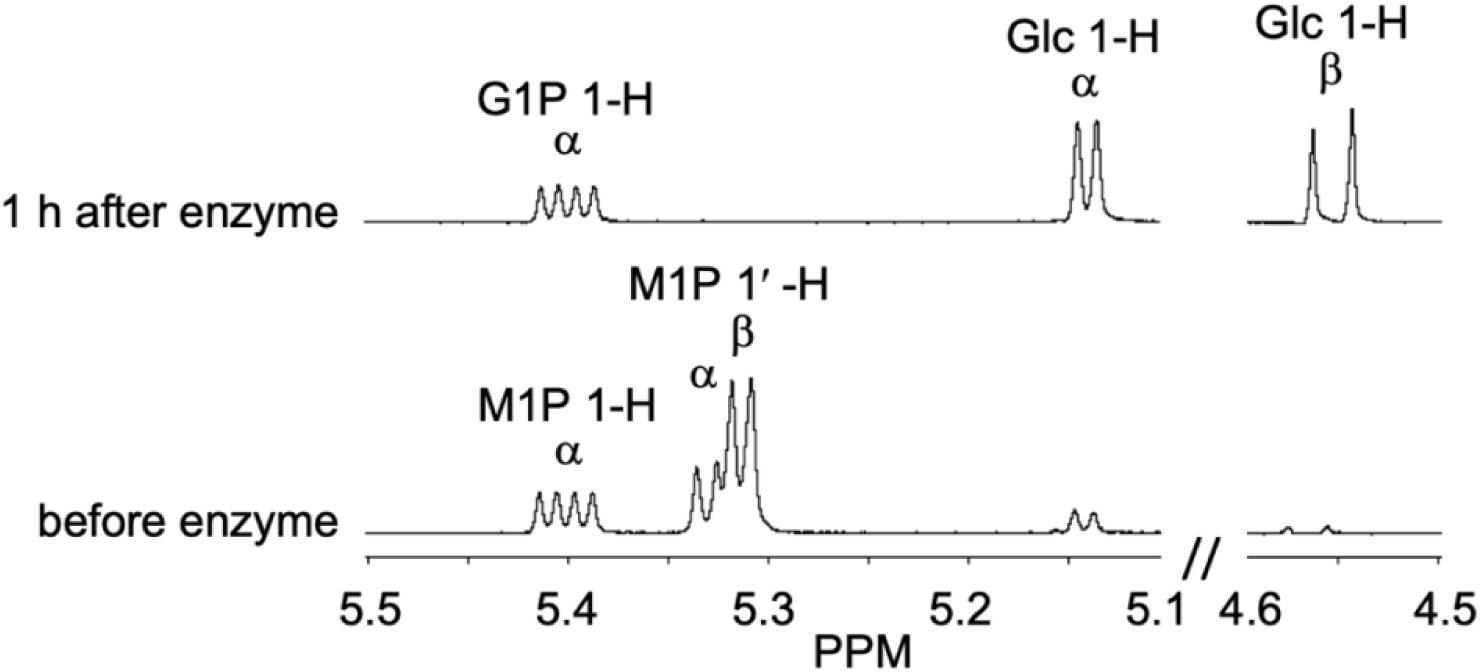
*A. niger* amyloglucosidase cleaves the α-1,4-linked non-reducing end glucose from both α and β-maltose-1-phosphate. An anomeric mixture of α/β-maltose-1-phosphate was incubated with amyloglucosidase. The reaction was monitored using ^1^H nuclear magnetic resonance spectroscopy and representative spectra are shown. The peaks for maltose-1-phosphate (M1P), glucose-1-phosphate (G1P) and glucose (Glc) were assigned as previously reported [10,23]. The 1′-H resonances of α/β-maltose-1-phosphate associated with the α-glucosyl-(1→4)-glucose linkage are lost at the same time as those of the 1-H reducing end of glucose appear. The 1-H resonances of α-maltose-1-phosphate and α-glucose-1-phosphate are essentially coincident. They are also slightly down-field compared with Fig. 3 because the solution was buffered at pH 4.0. Note that the 1-H resonances from β-maltose-1-phosphate and β-glucose-1-phosphate are not shown for clarity.

### 3.3. The H349Y GlgE mutation leads to a loss of enzyme activity and an increased temperature sensitivity

If the activity of GlgE is compromised by the H349Y mutation, it could manifest itself through a loss of either activity or stability. First, the kinetics of the extension of maltohexaose with α-maltose-1-phosphate by wild-type and H349Y GlgE were each monitored by the release of inorganic phosphate. It was immediately apparent that the activity of H349Y GlgE was lower than that of the wild-type enzyme. Nevertheless, the activity profiles of the two enzymes as a function of either pH or NaCl concentration were similar (Fig. 5A and 5B). Specifically, both enzymes showed optimal activity at pH 7.0, and NaCl up to 350 mM had little effect on either enzyme. Each enzyme had an acceptor preference for maltohexaose or maltoheptaose (Fig. 5C) with little or no difference between their acceptor profiles. The mutation therefore had little or no effect on these properties of GlgE despite a reduction in overall activity.

**Fig. 5.**
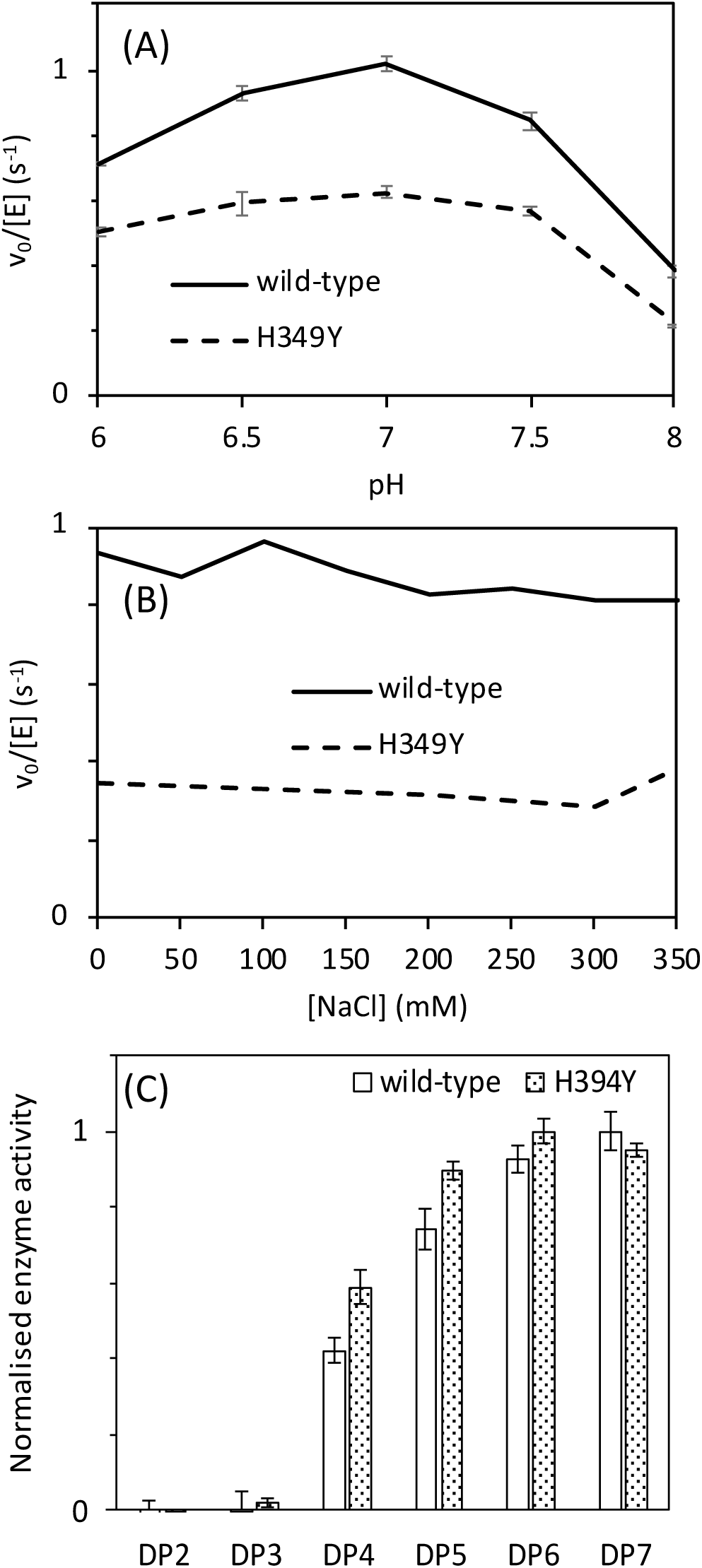
The H349Y mutation changes neither the pH optimum, salt-sensitivity nor acceptor specificity of *M. smegmatis* GlgE at 21 °C. Experiments were carried out using 0.25 mM α-maltose-1-phosphate and 1 mM acceptor. (A) shows the pH optima of wild-type (solid line) and H349Y (broken line) GlgE to be 7. Error bars represent standard errors (which are very small) from triplicate experiments. (B) shows the tolerance of wild-type (solid line) and H349Y (broken line) GlgE to NaCl up to 350 mM. (C) shows the acceptor preferences of wild-type (plain bars) and H349Y (shaded bars) GlgE are very similar. The acceptors tested ranged from maltose (degree of polymerisation (DP) of 2) up to maltoheptaose (DP of 7). Means are normalised to the highest value for each enzyme to aid comparison. The error bars represent standard errors from triplicate experiments.

To ascertain what effect temperature had on the wild-type and H349Y GlgE, their temperature stabilities were determined by pre-incubating each at a range of temperatures before assaying them at 21 °C. The wild-type GlgE enzyme showed a 24 to 23% drop in activity after exposure to temperatures of 40 and 45 °C, respectively (Fig. 6). By contrast, the activity of H349Y GlgE was more severely affected with drops in activity of 34 and 71% at 40 and 45 °C, respectively. From this trend, it is possible to estimate a 50% drop would have occurred at 42 °C. One can conclude that the H349Y mutation roughly doubled the rate of deactivation at 42 °C from ~24% to ~50% over a period of 20 min.

**Fig. 6.**
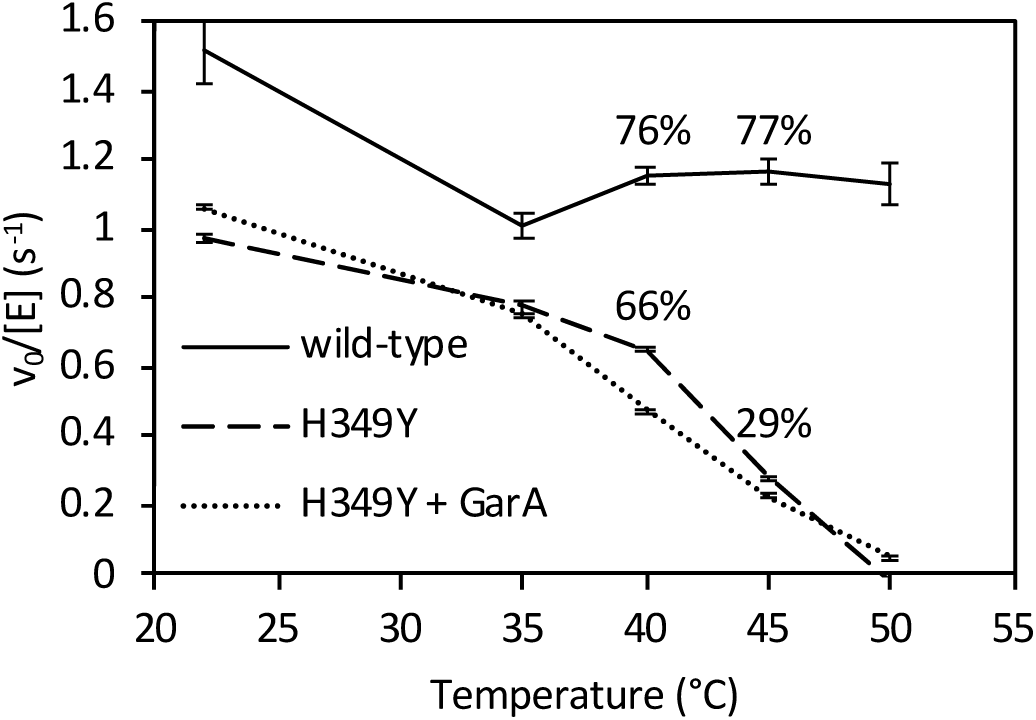
Temperature deactivation of wild-type and H349Y GlgE enzyme activity. Residual activity of wild-type (solid line) and H349Y GlgE in the absence (broken) or presence of GarA (dotted line) after pre-incubation at an elevated temperature for 20 min before assaying at 21 °C. Error bars represent standard errors of triplicate experiments. Percentage deactivation at 40 and 45 °C are indicated for wild-type and H349Y enzymes compared with their respective activities at 22 °C.

Over-expression of GarA *in vivo* resulted in a partial rescue of the temperature sensitive phenotype of SMEG53 cells [4] and GarA has been demonstrated to bind to and regulate not only phosphorylated but also non-phosphorylated targets [24]. To test whether GarA can directly protect H349Y GlgE from temperature de-activation, H349Y was pre-incubated with GarA. However, GarA showed no significant protective effect (Fig. 6). Indeed, it seemed to somewhat further destabilise H349Y GlgE.

The effect of temperature on GlgE activity was also determined. Enzyme activity was linearly dependent upon time over the 10 min period of the assay in all cases except for assays with the H349Y variant at 45 °C and above when initial rates had to be estimated. The wild-type enzyme showed an increase in activity between 25 and 45 °C, followed by a modest decrease at 50 and 55 °C (Fig. 7). By contrast, H349Y GlgE was considerably less active, and showed only a modest increase in activity between 25 and 40 °C, followed by a complete loss of activity at 50 °C and above, consistent with the temperature stability experiments. In contrast to the wild-type enzyme, there was no significant difference in the activity of the H349Y GlgE at 30 and 40 °C. Under these conditions, there were therefore ~3.5-fold and ~4-fold differences in turnover rates between the two enzymes at 30 and 40 °C, respectively.

**Fig. 7.**
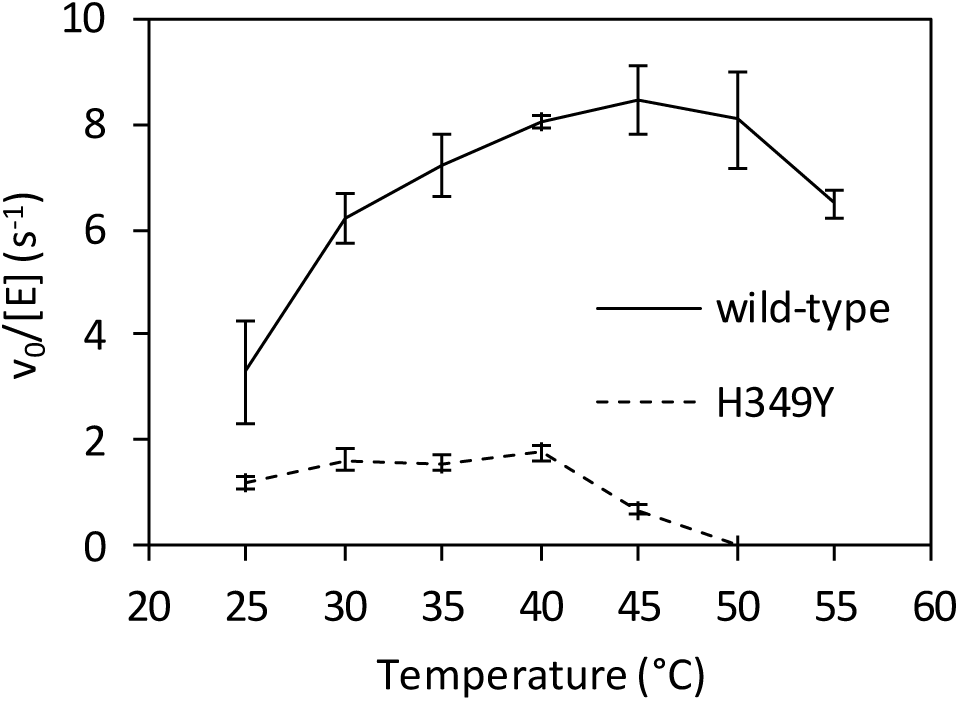
The effect of temperature on the enzyme activities of wild-type and H349Y GlgE. The activity of wild-type (solid line) and H349Y (broken) GlgE as a function of temperature was determined by monitoring phosphate release. Error bars represent standard errors of triplicate experiments.

To address whether the impaired activity of H349Y GlgE was due to a *k*_cat_ or *K*_m_ effect, the kinetic parameters with maltohexaose were measured for each enzyme at permissive and non-permissive temperatures for bacterial growth. Assays were conducted over a maximum of 10 min, minimising any impact of thermoinstability. The data exhibited substrate inhibition, so were fitted to a substrate inhibition model (Fig. 8 and Table 2). Substrate inhibition can be rationalised by high concentrations of maltohexaose promoting a competing disproportion reaction [1,10]. With the wild-type enzyme, both the *k*_cat_^app^ and *K*_m_^app^ for maltohexaose increased slightly at the higher temperature, resulting in a similar *k*_cat_^app^/*K*_m_^app^ value (Table 2).

**Table 2.** Kinetics of wild-type and H349Y GlgE. Enzyme activity was monitored by detecting phosphate release. Assays were performed in duplicate and values are expressed with standard errors. Quoted constants are apparent because the enzyme obeys ping-pong (substituted enzyme) kinetics [1]. See also Fig. 8.

| GlgE | T | $k_{cat}^{app}$ | $K_m^{app}$ | $K_i$ | $k_{cat}^{app}/K_m^{app}$ |
| --- | --- | --- | --- | --- | --- |
|  | °C | s<sup>-1</sup> | mM | mM | M<sup>-1</sup> s<sup>-1</sup> |
| wild-type | 30 | 78.9 ± 6.0 | 9.7 ± 1.5 | 126 ± 24 | 8100 ± 1400 |
| wild-type | 42 | 112 ± 14 | 14.4 ± 3.5 | 187 ± 67 | 7800 ± 2100 |
| H349Y | 30 | 15.0 ± 3.0 | 10.0 ± 3.8 | 95 ± 43 | 1500 ± 640 |
| H349Y | 42 | 14.5 ± 1.3 | 12.5 ± 2.3 | 366 ± 140 | 1160 ± 240 |

**Fig. 8.**
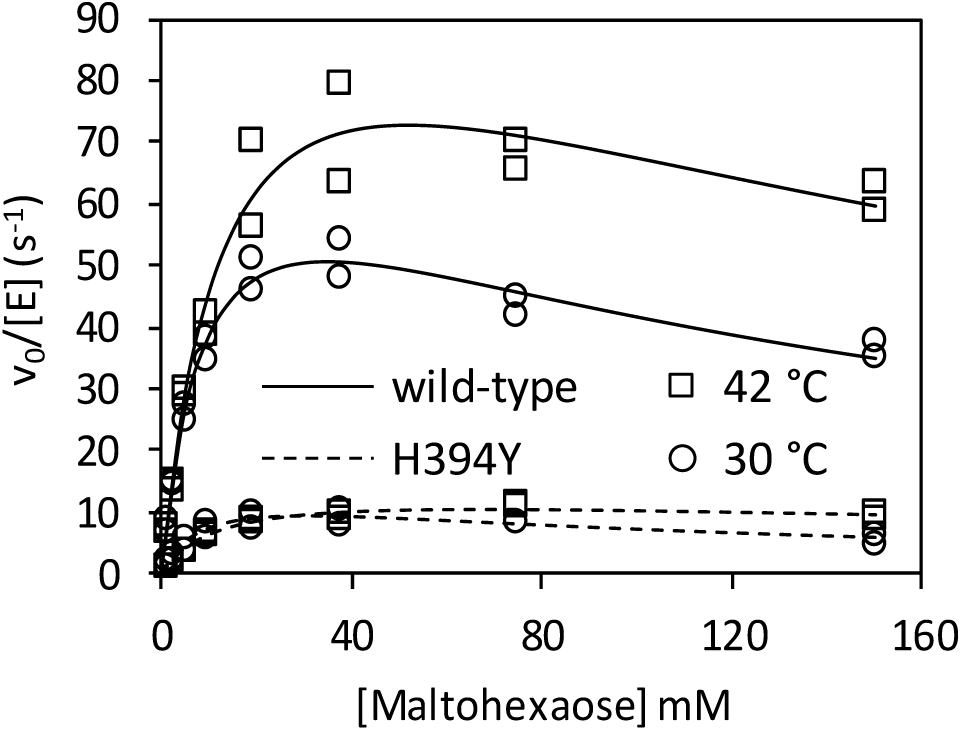
Kinetics of maltohexaose extension with α-maltose-1-phosphate by wild-type and H349Y GlgE. The kinetics were determined with enzyme concentrations between 12.5 and 100 nM and 1 mM α-maltose-1-phosphate at either 42 (squares) or 30 °C (circles) with wild-type (solid line) or H349Y (broken line). Experiments were carried out in duplicate. The lines of best fit adhere to a substrate inhibition model. See Table 2 for the derived kinetic constants.

The values of *K*_m_^app^ of H349Y GlgE for maltohexaose at each temperature were very similar to those of the wild-type enzyme. Given that *K*_m_^app^ is a function of the true *K* _m_ values for both substrates (equation 5), the affinity of neither substrate appears to have changed. By contrast, the values of *k*_cat_^app^, and therefore *k*_*cat*_^app^/*K*_*m*_ ^app^, were at five to seven-fold lower, consistent with the above experiments. Since *k*_cat_^app^ is a function of the true value of *k*_cat_, the H349Y mutation affects *k*_cat_ rather than the affinities of either substrate. Importantly, the *k*_cat_^app^/*K*_m_^app^ value for H349Y GlgE was about a third lower at the non-permissive vs the permissive temperature.

### 3.4. The H349Y mutation reduces thermal stability but has limited impact at the non-permissive temperature

Circular dichroism spectroscopy was used to probe the secondary structure and temperature stability of wild-type and H349Y GlgE. The two forms of GlgE had very similar net secondary structure at 25 and 42 °C (Fig. 9 and Table 3). After 2 h of incubation at 42 °C, the H349Y enzyme showed a shift in the spectrum compared with the wild-type enzyme (Fig. 9). However, this shift was very small, equating to changes in secondary structural elements of ≤ 2% each (Table 3). Melting temperatures were determined by monitoring the loss of secondary structure as a function of temperature. The H349Y mutation led to a 7.6 °C destabilisation of the GlgE protein from 55.7 ± 0.2 to 48.1 ± 0.3 °C. Consistent with this observation, a Thermofluor assay that follows the thermal denaturation of proteins via binding of a fluorescent dye was in close agreement, showing a 9 °C destabilisation from 56 to 47 °C (Fig. 10). At 42 °C, the amount of melting of the wild-type and H349Y enzymes was 1 and 16%, respectively. Therefore, no gross denaturation of the protein occurred at the non-permissive temperature for bacterial growth, consistent with only partial loss of activity for the H349Y enzyme at 42 °C (Fig. 6).

**Table 3.** Net secondary structures of wild-type and H349Y *M. smegmatis* GlgE. The values were determined using circular dichroism spectroscopy (see Fig. 9). Values are means of triplicate measurements. Note, the elements add up to either 99, 100 or 101 depending on rounding introduced by the software.

| GlgE | T | t | % helix | % sheet | % turns | % unstructured |
| --- | --- | --- | --- | --- | --- | --- |
|  | °C | h |  |  |  |  |
| wild-type | 25 | 0 | 22 | 30 | 21 | 28 |
| wild-type | 42 | 0 | 21 | 28 | 20 | 30 |
| wild-type | 42 | 2 | 20 | 32 | 20 | 29 |
| H349Y | 25 | 0 | 23 | 26 | 20 | 31 |
| H349Y | 42 | 0 | 20 | 31 | 20 | 29 |
| H349Y | 42 | 2 | 18 | 30 | 21 | 30 |

**Fig. 9.**
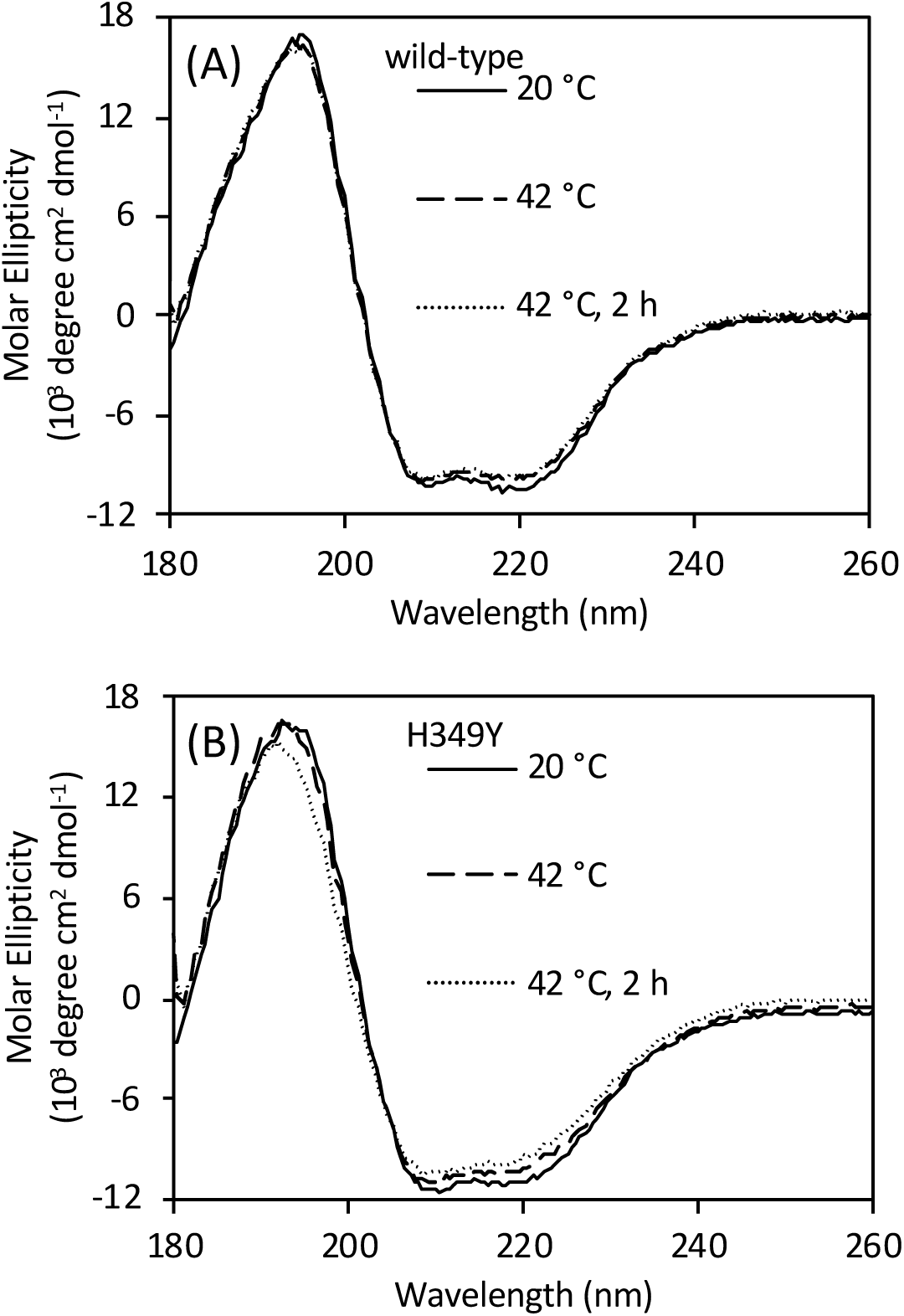
Circular dichroism spectra of wild-type (panel A) and H349Y (panel B) GlgE. Spectra were recorded at either 20 or 42 °C, as indicated. An additional spectrum was recorded for each enzyme after 2 h of incubation at 42 °C. See Table 3 for the corresponding analysis of net secondary structures.

**Fig. 10.**
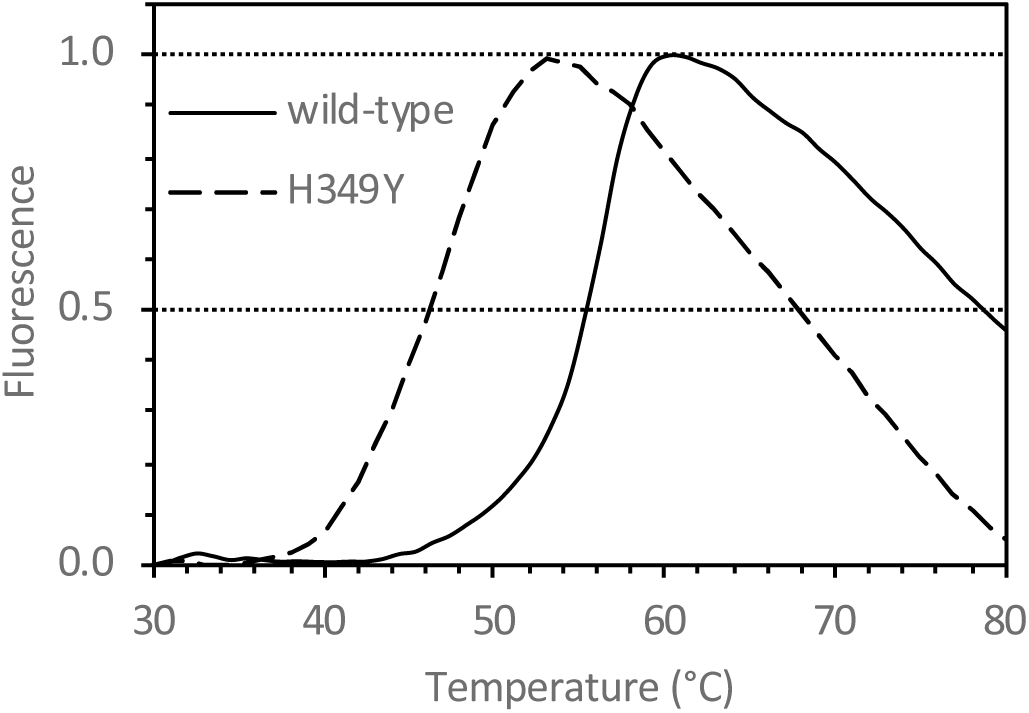
Thermofluor analysis of wild-type and H349Y GlgE. The proteins were each heated in the presence of CYPRO Orange. Fluorescence (arbitrary units) is shown normalised to the maximum signal of each sample. The melting temperatures for wild-type (solid line) and H349Y (broken line) GlgE were 56 and 47 °C, respectively, as highlighted by the horizontal guideline at 0.5. The normalised fluorescence values at 42 °C for wild-type and H349Y GlgE were 0.01 and 0.16, respectively.

### 3.5. Neither GlgE nor phospho-GlgE bind to GarA

Although GarA was shown above not to rescue the temperature sensitivity of the H349Y mutation, it was not possible to rule out a protein-protein interaction between the two. We therefore employed several methods to explore whether any interaction could occur. A surface plasmon resonance approach was not possible because neither protein bound successfully to a CM5 sensor chip at all values of pH tested (4.0 and 4.5). Similarly, only non-specific binding was observed when attempting to use an NTA sensor chip to immobilise each His-tagged protein. We therefore took another approach. GarA has no tryptophan residues. By contrast, GlgE has intrinsic fluorescence with its 17 tryptophan residues, which could change when another protein binds to GlgE. However, the intrinsic fluorescence of GlgE (at 1 µM) did not change in the presence of GarA (up to 100 µM). In separate experiments, the labelling of GlgE with dansyl groups allowed the measurement of fluorescence anisotropy. This signal would change when the tumbling rate of GlgE slows on binding another protein. However, no such change was detected in the presence of GarA. It was not possible to repeat the experiment with unlabelled GlgE and labelled GarA due to protein solubility limitations. Nevertheless, given the sensitivity of fluorescence anisotropy, one can conclude that GlgE and GarA do not interact in the conditions tested.

## 4. Discussion

We have shown that the temperature-sensitive mutation in the *glgE* gene leads to the accumulation of α-maltose-1-phosphate at the non-permissive temperature (Table 1 and Fig. 3). This is entirely consistent with the phenotype of other mutations in mycobacterial *glgE* reported recently [1,5], where a loss of GlgE activity leads to the accumulation of α-maltose-1-phosphate and the slowing of bacterial growth. However, this contrasts with the 1999 study [4], where it was concluded that the accumulation of glycogen led to the cessation of bacterial growth. It is easy to reinterpret the 1999 study with hindsight, but the conclusions drawn at the time were entirely reasonable. First, GlgE was predicted to be a glucanase and second, the only known potential precursors of glycogen at that time were UDP-glucose or ADP-glucose, which would not have been susceptible to hydrolysis by amyloglucosidase. By contrast, the recently discovered α-maltose-1-phosphate precursor possesses a glucosyl-(1→4)-glucose linkage that is susceptible to hydrolysis, leading to the liberation of glucose. Since the standard glycogen assay involves the detection of glucose liberated by amyloglucosidase, α-maltose-1-phosphate would have given a false positive result. It therefore follows that the interpretation of the observations in 1999 that led to the conclusion that glycogen recycling was essential for growth were based on an assumption that we now know to be incorrect. It therefore remains to be seen whether glycogen recycling is indeed essential for growth.

The basis for the temperature-sensitivity of the H349Y mutation appears not to be a simple case of protein instability at the non-permissive temperature. Relative to the wild-type enzyme, the H349Y enzyme lost only ~half of its activity after 20 min of incubation at 42 °C (Fig. 6) with only minimal changes in net secondary structure after 2 h (Fig. 9 and Table 3) and only 15% more melting (Fig. 10). Rather, it appears to be also down to the loss of activity of the enzyme at the non-permissive temperature caused by a ~7-fold reduction in the *k*_cat_^app^ relative to the wild-type enzyme (Fig. 8 and Table 2). This was due to a much smaller intrinsic value of *k*_cat_^app^ (Fig. 8 and Table 2) coupled with a much smaller apparent temperature coefficient than the wild-type enzyme (Fig. 7) leading to a smaller *k*_cat_^app^ for the H349Y variant at the higher vs lower temperature. Since there was a ~5-fold reduction in activity caused by the mutation at the permissive temperature (Fig. 8), this implies that adequate flux through the GlgE pathway is on a knife-edge somewhere between the permissive and non-permissive temperatures. At the higher temperature, the accumulation of α-maltose-1-phosphate is evidently correlated with reduced bacterial growth-rate, consistent with all previous studies [1,5].

The H349Y mutation of GlgE maps onto the surface of the protein (Fig. 11) that is on the opposite side of the molecule to the substrate binding sites [10,25]. The closest residue that forms part of the substrate binding sites is within five amino acids within the sequence. Note that this His residue is highly conserved in GlgE enzyme across mycobacteria, including *M. tuberculosis*, and streptomycetes. Quite how the substitution affects enzyme activity is therefore not clear, which mirrors our lack of understanding of how phosphorylation lowers activity [14].

**Fig. 11.**
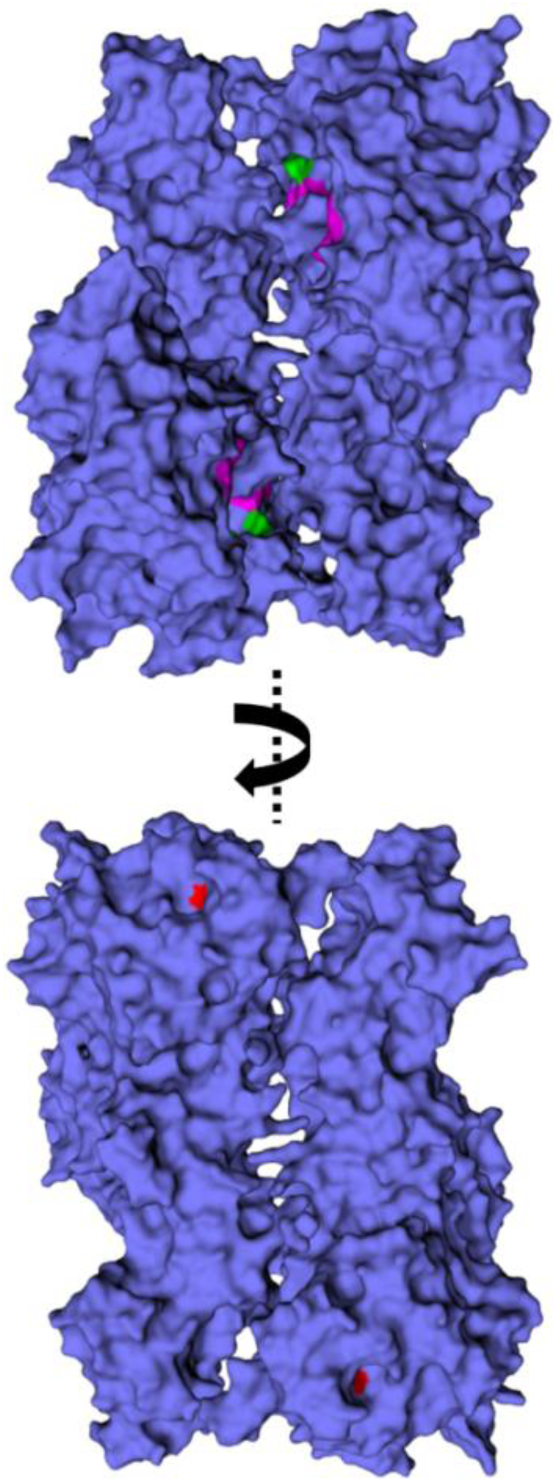
The location of the substituted His residue on GlgE. The structure of *Streptomyces coelicolor* GlgE isoform I is shown with surface representation (PDB entry 5LGV) [25]. This structure is of a Glu423Ala variant with a maltooligosaccharide bound to both the donor and acceptors sites (with this ligand omitted for clarity). The equivalent residue to His349 in the *M. smegmatis* enzyme is His329 in the *S. coelicolor* enzyme, which is highlighted in red and is visible in the lower panel. Representative residues associated with the binding site of α-maltose-1-phosphate (catalytic residues Glu423Ala and Asp394, together with Gln324 and Asn352; *S. coelicolor* numbering) are highlighted in green, while some of those associated with the binding site of the acceptor (His397, Trp448 and Arg449) are highlighted in magenta. All of these residues associated with the binding of substrates are visible in the upper panel.

GarA has been reported to rescue the temperature-sensitive mutation by reducing the accumulation of carbohydrate [4]. All attempts to identify an interaction between GarA and GlgE suggested no such interaction occurs. In addition, GarA did not strongly influence the temperature stability of GlgE (Fig. 6). Both GarA and GlgE are substrates of the phosphorylase PknB. Phosphorylation of GarA precludes its binding to other proteins. Phosphorylation of GlgE leads to its negative regulation, primarily through a lowering of *k*_cat_^app^ by up to two orders of magnitude [14]. Perhaps GarA binds to phosphorylated GlgE and stabilises it. However, preliminary anisotropy experiments with GarA and GlgE phosphorylated by PknB indicated no interaction (data not shown). In any case, phosphorylated GlgE has very low activity, so its stabilisation would not recover flux through the enzyme. The most likely explanation for the ability of GarA to rescue the mutation is therefore that it acts as a decoy at high concentrations, preventing the negative regulation of GlgE. It therefore follows that GlgE would normally be somewhat negatively regulated by PknB at the non-permissive temperature, perhaps to counter its temperature-dependent increase in activity (Fig. 7). It could therefore be the combination of negative regulation, thermo-instability and a reduced *k*_cat_^app^ that breaks the threshold for tolerance of α-maltose-1-phosphate toxicity in the mutant at the higher temperature leading to a lower rate of bacterial growth.

## 5. Conclusions

In this study, we have shown that it is possible to reconcile the original report that GlgE is a glucanase [4] with what is now known about the biosynthesis of glycogen in mycobacteria [5]. When GlgE is compromised by the H349Y amino acid substitution, α-maltose-1-phosphate accumulates to toxic levels due to a partial loss of polymerase activity, particularly at an elevated temperature, coupled with a loss of stability. This implies the regulation of GlgE is delicately balanced to control flux through the glycogen biosynthetic pathway without compromising growth. This increases the attractiveness of the GlgE pathway as a potential drug target [26].

## Author Contributions

KS carried out the enzymology. SFDB carried out the bacterial growths and metabolite analyses. SS and RK constructed the microbial strains and also carried out bacterial growths. SB and RK conceived the project and supervised the work. SB wrote the manuscript from a draft written by KS with subsequent feedback from all authors.

## Declaration of Competing Interest

The authors declare that no conflict of interest exists.

## Acknowledgements

We thank Virginie Molle for the *garA* and *pknB* expression vectors and for feedback on the manuscript.

## Funding

This work was supported by the United Kingdom Biotechnology and Biological Sciences Research Council (Doctoral Training Partnership [BB/J014524/1] and Institute Strategic Programme [BB/J004561/1] grants) and the John Innes Foundation. R.K acknowledges support from the Jürgen Manchot Foundation.

## Notes

### Competing Interest Statement

The authors have declared no competing interest.

### Summary of Updates

New data have been included and clarifications have been made throughout. The conclusion has been revised such that enzyme thermo-instability of GlgE contributes to the temperature-sensitive phenotype along with a lower intrinsic turnover number for the enzyme.

